# Environmental enrichment on neural plasticity in juvenile Baltic sturgeon (*Acipenser oxyrinchus*)

**DOI:** 10.1101/491910

**Authors:** Maria Cámara-Ruiz, Carlos Espirito Santo, Jörn Gessner, Sven Wuertz

**Author notes:** **Corresponding author**: M. Cámara-Ruiz.

## Abstract

Conservation biologists have long emphasized the importance of environmental enrichment in hatcheries which main objective is restocking. In this study, we evaluated the impact that long-term exposure of environmental enrichment in an artificial river section (surface water, ambient light and temperature) has on neural plasticity in juvenile Baltic sturgeons (*Acipenser oxyrinchus*) in comparison to fish that were reared in glass-fiber tanks in a recirculation system (aquaculture control group) under constant light and temperature conditions and an intermediate group reared in glass-fiber tanks with ambient water and natural photoperiod and temperature fluctuation. All expositions were carried out in triplicate. From the three markers reported to indicate neural plasticity, only an up-regulation of *neurod1* in the forebrain of Baltic sturgeon (*A. oxyrinchus*) reared in enriched environment was observed in comparison to the semi-natural group and to the aquaculture (control) group. The two other markers (brain-derived neurotropic factor (*bdnf*), and proliferating cell nuclear antigen (*pcna*) did not reveal any significant differences between the groups. In conclusion, only minor effects of environmental enrichment on neuroplasticity were observed here.

## 1. Introduction

During the last 150 years, sturgeons (Acipenseridae) have experienced a drastic decline due to several reasons including among others overfishing, habitat destruction and pollution. In fact, sturgeons are considered one of the most endangered fish species worldwide (IUCN, 2018). Baltic sturgeon (*Acipenser oxyrinchus*), is currently considered extinct or missing in the range countries to the Baltic Sea. In order to recover Baltic sturgeon populations, restoration programs have been initiated in an attempt to establish self-sustaining populations and thus to safeguard the long-term viability of the species (Gessner et al., 2011).

Several authors consider that sturgeon stock enhancement contributes significantly to sustain populations as it has been seen in the Caspian Sea, Azov Sea or in the Volga River (Billard & Lecointre, 2001). However, there are still major issues regarding the survival of hatchery-reared fish during the first days post-release. One of the impacts that reduce the overall success of the fish released is associated to maladaptation of stocking material reared under conventional hatchery conditions. Inability to obtain feed, aggressiveness, lack of predator recognition and a reduced brain size in general are among the reasons for the poor performance. The reason for such poor results may be due to the classical hatchery environments being deficient of structure and lacking appropriate stimuli, resulting in fish not fit to survive in the natural habitat (Miller et al., 2004; Salvanes et al., 2007; Johnsson et al., 2014). It was suggested that environmental enrichment would be an appropriate means to circumvent the problems listed above and to increase fitness and resultantly survival upon release (Berejikian et al., 2001; Brown et al., 2003; Salvanes et al., 2013).

Environmental enrichment has come to different meanings in the literature; for instance, increasing the structural complexity of the environment or providing a first approach of the habitat the animal is likely to be exposed to in the wild could be considered as environmental enrichment (Sheperdson, 1994). However, conservation biologists have emphasized the importance of practices such as environmental enrichment, pre-release training or soft release strategies in restoration programs. To date, the production of ecologically viable individuals is mostly not taken into consideration because hatchery operations focus on the production of large quantities of fish, rather than on the fitness of the fish.

In their natural habitats, fishes experience environmental challenges and they have to adapt their physiology and behavior in order to physiologically and behaviourally adjust to such changes. Much of this flexibility is supported and influenced by cognition and neural plasticity (Ebbesson and Braithwaite, 2012). Thus, changes within the brain (structural or neurophysiological) are believed to lead to differences in behavioral phenotypes (Salvanes et al., 2013). In general, captive-bred animals have reduced behavioral flexibility and many regions of the brain are smaller and less active in comparison to wild conspecifics (Salvanes et al., 2013), which appears to be a result of the non-demanding hatchery rearing environment.

Neuroscientists have studied the molecular mechanisms of neural plasticity associated with memory. This work has resulted in markers related to neural plasticity. Recent studies have indicated that pro-neural gene neurogenic differentiation 1 factor (*neurod1*) is a reliable measure of neurogenesis in fish and a useful indicator of the neural plastic changes associated with memory and learning (Rossi et al.,2006; Grassie et al., 2013). Moreover, brain-derived neurotrophic factor (*bdnf*) has an important role in neural plasticity through sculpting and refinement of synapses and through promoting neurogenesis and cell survival (Castrén and Rantamäki, 2009). Although not specific to the brain, proliferating cell nuclear antigen (*pcna*) is a marker for cell proliferation in the respective organ (Leung et al., 2005). Taking into account the functions of the genes previously mentioned (*neurod1*, *bdnf* and *pcna*), they were of interest for this present study.

Preliminary work in which hatchery-reared fish were exposed to outside ponds before release suggests that this brief exposure to a natural environment significantly improves survival rates (Maynard & Flagg, 1994). Taking this into account, the main objective of this study was to evaluate the impact that long-term rearing of juvenile Baltic sturgeons, *A. oxyrinchus,* in an environmentally enriched pond mimicking a river stretch has on neural plasticity when compared to conventional and semi-.natural rearing environments.

## 2. Materials and methods

### 2.1. Experimental design

The experiments were performed at the experimental facilities at the Leibniz Institute of Freshwater Ecology and Inland Fisheries (IGB, Berlin, Germany) using fish from the sturgeon stock kept at the IGB. Juvenile Baltic sturgeons (*Acipenser oxyrinchus*) were reared in different environments (aquaculture, semi-natural and natural) over a period of 6 months, each group assessed in triplicate. Fish were fed twice a day with chironomids at 5% of the body weight. 180 fish were randomly distributed to the triplicates of each group (20 fish per tank). The aquaculture control group was reared at constant light (12:12L:D) and temperature (20±1,2°C) regime in 1.2×1.2×0.44m green glass fiber tanks following classical aquaculture protocols The semi-natural group was reared in green glass fiber tanks (1×1×0.6m) outside, exposed to natural temperature and light conditions while the natural group was reared in artificial river stretches installed in a pond, exposed to natural temperature and light conditions and also to a more complex environment.

For the assessment of brain plasticity, fish from the three groups were sampled after 6 months of experimental trial. Fish were euthanized with MS222 (300 ppm) followed by cutting through the spinal cord. Brains from 27 Baltic sturgeons (n=9) were dissected and divided into three parts representing the three main brain regions (forebrain, midbrain and hindbrain). Samples were stored in RNA later at −80 °C for subsequent gene expression analysis.

### 2.2. Gene expression

Total RNA was extracted with TRIzol as described by Reiser et al. (2011), including a DNase I digestion. Total RNA concentration and purity were determined in duplicates with a Nanodrop® ND-1000 UV–Vis spectrophotometer. Purity was validated as the ratio of the absorbance at 260 and 280 nm (A260/280) ranging between 1.8 and 2.0. Moreover, integrity of the total RNA was checked by gel electrophoresis and, in 10% of all samples, on RNA 6000 Nano chips with an Agilent 2100 Bioanalyzer. To eliminate potential DNA contamination, DNAse I digestion was performed in all samples prior to transcription. Next, mRNA was transcribed with MMLV Affinity reverse transcriptase (Agilent, 200 Units/μl) according to the manufacturer’s instruction. In 10% of the samples, the enzyme was substituted by pure H_2_0, serving as a control (-RT) to monitor DNA contamination.

Species-specific primers targeting elongation factor 1α (*ef1a*), brain-derived neurotropic factor (*bdnf*), neurogenic differentiator factor (*neurod1*) and proliferating cell nuclear antigen (*pcna*) were designed using the sequence information available. Specificity of the assays was confirmed by direct sequencing (SeqLab, Germany). Real-time PCR was carried out with Mx3005p qPCR Cycler (Stratagene), monitoring specificity by melting curve analysis. Full specifications of qPCR assays, including primer sequences are given in Table 1.

**Table 1.**
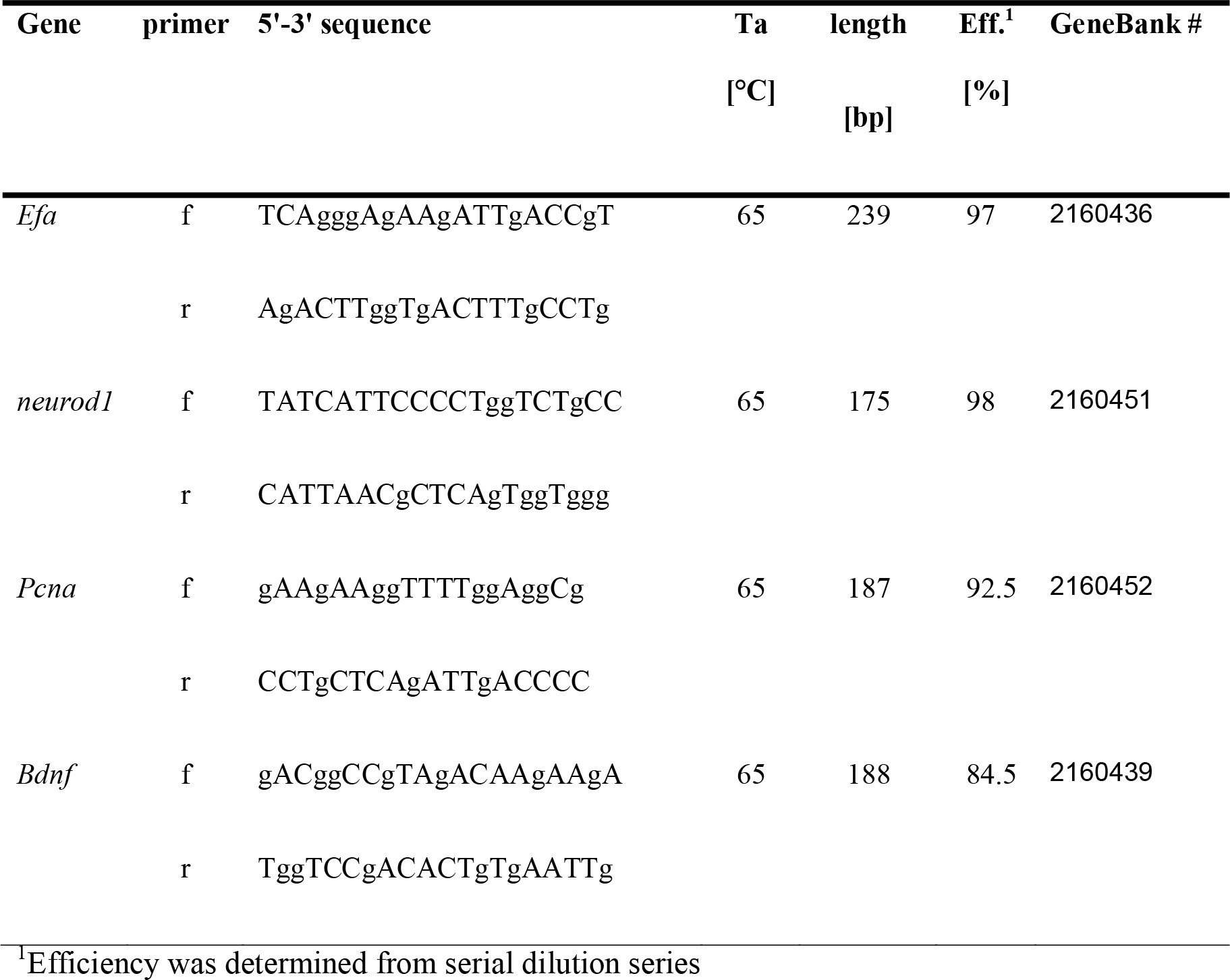
Specifications of qPCR assays including primer sequences, annealing temperature (Ta), amplicon length [bp], PCR efficiency (Eff) and NCBI accession number of the respective housekeeping (ef) and target genes: ef-elongation factor 1 a, neurod1-neurogenic differentiaton factor, bdnf - brain-derived neurotrophic factor, pcna-proliferating cell nuclear antigen-

For the RT-QPCR, 2 μL of the diluted sample (40 ng/μL) were used as template in 20 μL PCR mix [SYBR-Green I (Invitrogen), 200 μM of each dNTPs (Qbiogene), 3 mM MgCl 2 and 1 U Invitrogen Platinum Taq polymerase]. PCR conditions comprised an initial denaturation at 96 °C for 3 min, followed by 40 cycles of denaturation at 96 °C for 30 s, primer annealing (for Ta, see Table 1) for 30 s and elongation at 72 °C for 30 s. PCR efficiencies were determined experimentally with a dilution series of a calibrator corresponding to 200 ng/μl. PCR assays for all individual samples were run in duplicate. Expression of target genes were calculated by the comparative CT method (ΔΔCT) according to (Pfaffl, 2001), correcting for the assay efficiencies and normalizing to elongation factor 1α (*ef1a*) as a housekeeping gene. Expression data are presented as fold increase.

### 2.3. Data analysis and statistical methods

Data are presented as mean ± standard deviation (SD). Prior to statistical analyses, all data were tested for normality of distribution using the Kolmogorov-Smirnov/ Shapiro Wilk test and for homogeneity using Levene’s test. Data on RNA expression was analyzed with a parametric Tukey’s test or non-parametric Dunn’ test. The level of significance used was P ≤ 0.05. All statistical analyses were performed with GraphPad Prism statistical program.

## 3. Results

### 3.1. Brain plasticity and cognition

Selected genes related to brain plasticity and cognition (*bdnf*, *neurod1*, *pcna*) were analyzed in all three brain areas of Baltic sturgeon at the end of the rearing trial. Regarding the forebrain region (Fig. 1), significant differences were observed only in the expression of *neurod1*. In particular, there was an up-regulation in the natural group in comparison to the semi-natural group in Baltic sturgeon after 6 months of experimental trial (Fig. 1). The differences determined between the natural and the aquaculture/control groupwere recognizable but not significantly different in the gene expression of *neurod1*. No significant differences were observed between the semi-natural and the aquaculture/control group (Fig.1). Also, the gene expression of *pcna* and *bdnf* in the forebrain was not significantly different. No significant differences were observed in the expression of the selected genes for the midbrain and hindbrain either (Figs 2 & 3).

**Figure 1.**
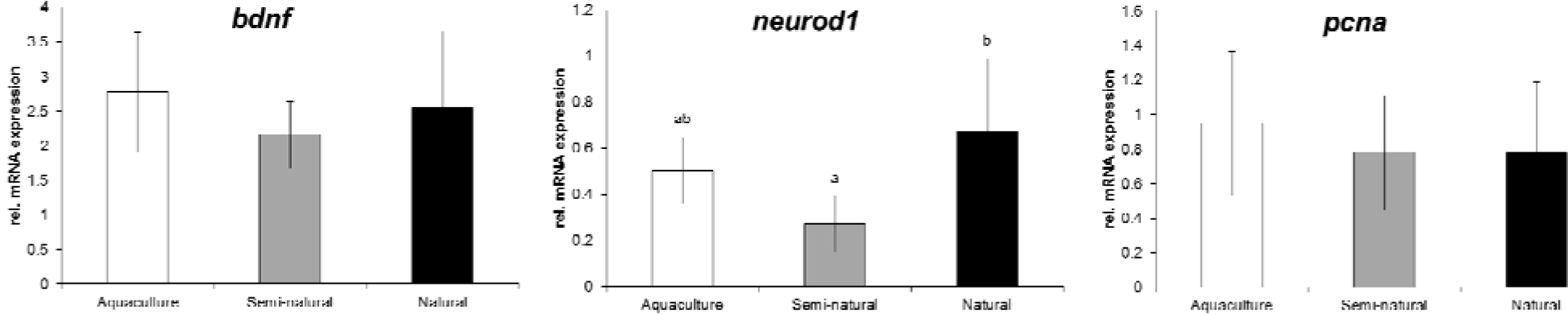
Gene expression, determined by qPCR, in the forebrain of Baltic sturgeon (*A. oxyrinchus*) of the aquaculture, semi-natural and natural groups after 6 months of experimental trial. Data are expressed as fold relative. Groups with different subscripts are significantly different (P < 0.05, n = 9). Statistic test used (Dunn’s or Tukey’s) is indicated below the respective graph.

**Figure 2.**
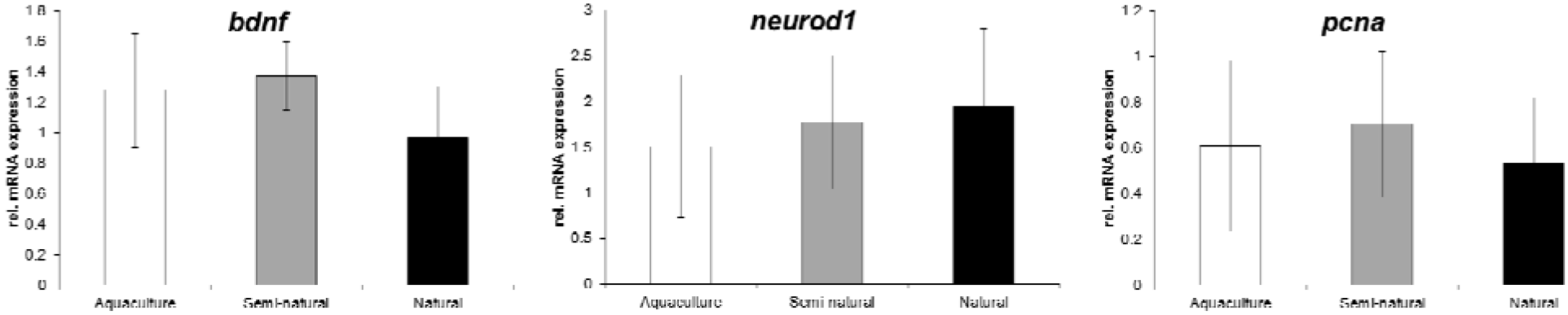
Gene expression, determined by qPCR, in the midbrain of Baltic sturgeon (*A. oxyrinchus*) of the aquaculture, semi-natural and natural groups after 6 months of experimental trial. Data are expressed as fold relative. Groups with different subscripts are significantly different (P < 0.05, n = 9). Statistic test used (Dunn’s or Tukey’s) is indicated below the respective graph.

**Figure 3.**
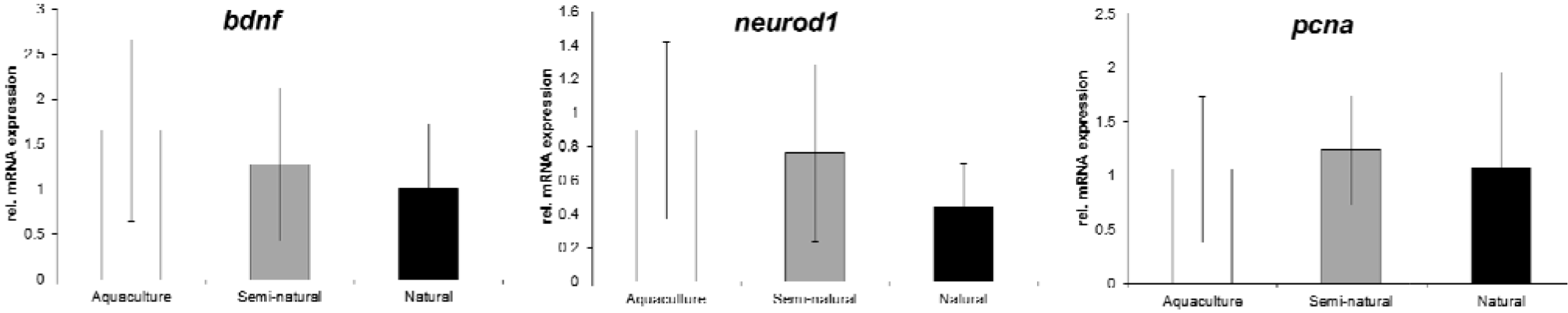
Gene expression, determined by qPCR, in the hindbrain of Baltic sturgeon (*A. oxyrinchus*) of the aquaculture, semi-natural and natural groups after 6 months of experimental trial. Data are expressed as fold relative. Groups with different subscripts are significantly different (P < 0.05, n = 9). Statistic test used (Dunn’s or Tukey’s) is indicated below the respective graph.

## 4. Discussion

It is known that controlled rearing environments are homogeneous and impoverished in comparison to natural environments since the variability of abiotic and biotic factors is generally much lower in the hatchery than in the wild. As a result, hatchery-reared fish are predicted to be less able to properly adjust to the full range of variation in the natural environment when released than their wild conspecifics (Johnsson et al., 2014). A potential solution to this problem could be an increase of environmental variability during the rearing period. In fact, many of the captive-breeding programs for mammals allow the animals to experience natural environments or alternative habitats that contain naturalistic features of the environment into which the animals are more likely to be released afterwards (Sepherdson et al. 1994).

Over the last years, there has been growing attention in the use of environmental enrichment to promote behavioural flexibility in animals that are bred for release, specifically in fish (Berejikian et al., 2001; Brown et al., 2003; Strand et al., 2010). Within the brain, environmental enrichment has been shown to affect neurogenesis and synaptic plasticity in the hippocampus (Salvanes et al., 2013). There is growing (but not always consistent) evidence that the addition of enrichment increases behavioural flexibility and researchers have started to investigate the effects of enrichment on fish brain. Recent studies have shown that expression levels of the pro-neuronal gene neurogenic differentiation 1 (*neurod1*) are a reliable measure of neurogenesis and a useful indicator for neurogenic changes (Rossi et al., 2006; Grassie et al., 2013).

In the present study, an up-regulation in the pro-neuronal gene neurogenic differentiation 1 (*neurod1*) in the forebrain of Baltic sturgeon (*A. oxyrinchus*) reared under nature-like conditions for a period of 6 months was observed in comparison to the group reared under semi-natural/intermediate conditions. We also observed a tendency of an up-regulation in the expression of *neurod1* in the forebrain of Baltic sturgeon from the enrichment group in comparison to the aquaculture (control) group. Thus, our results only moderately support the hypothesis that enrichment during the hatchery period had positive effects on neural plasticity in Baltic sturgeon. The insignificant differences in expression of the *bdnf* and *pcna* could indicate that the difference in stimuli was not strong enough to result in substantial differences or that long-term adaptation results in homogenization of the expression. These results are in agreement with previous studies such as Salvanes (2013), which found an up-regulation in the forebrain expression of *neurod1* after exposing Atlantic salmon (*Salmo salar*) environment enrichment and an improvement in learning ability. Moreover, another study in Atlantic cod (*Gadus morhua*), experience with spatial structures in the nursery environment increased behavioral flexibility (Salvanes & Braithwaite 2005; Salvanes et al. 2007).

Furthermore, previous studies have addressed the effect of enriched environments in fish and they have reported a range of behavioral benefits such as adaptations in foraging abilities, decreased aggression or improved social learning skills between others (Salvanes 2005; Berejikian 2001). All these adaptations together are likely to result in an increase of survival post-release. If hatchery-reared fish manage to survive the first week after release, then the chance of long-term survival is greatly increased (Kanidhev et al 1970, Brown and Smith 1998). Since environmental enrichment in captive environment increases behavioural plasticity it could potentially increase the chance of survival upon release. As such it would be a sensible measure to be considered in conservation restocking where quality is more important than quantity.

## Acknowledgments

The authors thank Eva Kreuz for her contribution in the laboratory. The study was supported by the European Training Network of the Marie Sklodowska-Curie Actions ITN “Improved production strategies of endangered freshwater species. This project has received funding from the European Union´s Horizon 2020 research and innovation program under the Marie Sklodowska-Curie grant No [642893].

## Competing Interests

I declare that there are no competing interests.

## Author contributions

The experiment was conducted by S.W. and C.E.S.. The laboratory analysis was carried out by M.C.R. M.C.R. wrote the first draft of the manuscript. S.W. supervised the project. The manuscript was revised by all co-authors.

